# The extension of mammalian pregnancy required taming inflammation; independent evolution of extended placentation in the tammar wallaby

**DOI:** 10.1101/2023.06.18.545206

**Authors:** Jessica S. Dudley, Marilyn B. Renfree, Günter P. Wagner, Oliver W. Griffith

**Affiliations:** School of Natural Sciences, Faculty of Science and Engineering, Macquarie University, NSW, 2109, Australia; School of BioSciences, University of Melbourne, Melbourne, Victoria, 3010, Australia; Department of Ecology and Evolutionary Biology, Yale University, New Haven, CT, 06520, USA; Yale Systems Biology Institute, Yale University, West Haven, CT, 06520, USA; Department of Obstetrics, Gynecology and Reproductive Sciences, Yale Medical School, New Haven, CT, 06520, USA

**Keywords:** marsupial, placenta, implantation, recognition of pregnancy, gene expression, inflammation

## Abstract

In the first live bearing mammals, it is assumed that pregnancy was short and ended with a brief period of inflammatory maternal-fetal interaction. This mode of reproduction has been retained in many marsupials. While inflammation is key to successful implantation in eutherians, a key innovation in eutherians is the ability to switch off this inflammation after it has been initiated. This extended period, in which inflammation is suppressed, likely allowed for an extended period of placentation. One lineage of marsupials, the macropodids (wallabies and kangaroos), have extended placentation beyond the 2-4 days seen in other marsupial taxa, which allows us to test whether a moderated inflammation response after attachment is a general pattern associated with the extension of placentation in mammals. We show that during tammar wallaby pregnancy, some inflammatory genes are expressed at key time points of gestation, including *IL6*, before attachment, *IL12A* and *LIF* throughout the period of placentation and prostaglandins before birth. However, we did not see evidence of a complete inflammatory response at any time point. We argue that genes involved in a moderated inflammation reaction may have been co-opted into roles for placentation, facilitating the establishment and maintenance of extended fetal-maternal contact. Whilst the absence of other key mediators of inflammation may prevent prolonged damage to the uterus. We argue the moderation of inflammation following maternal-fetal contact is a convergently evolved key innovation that allowed for the extension of placentation in different mammalian lineages.

**Significance statement:** Our data suggest that moderation of the inflammatory reaction to embryo attachment allows for extension of pregnancy in mammals. The ancestor of all mammals likely experienced an ancestral inflammatory reaction in response to embryo attachment. In contrast, eutherians and some marsupials, such as macropodids, have an extended period of fetal-maternal contact. During this period of placentation many inflammatory genes are silenced while a few others are still expressed. This moderated expression of inflammatory genes suggests that some genes of inflammation were coopted into establishing and maintaining the placenta. This challenges the perspective of inflammation as being detrimental to pregnancy, instead suggesting that fetal-maternal interactions are based on a modified inflammation response necessary for maintaining pregnancy over an extensive period of time.

## Introduction

In the first live-bearing mammals, pregnancy was short with only a brief period of maternal-fetal attachment (1). This period of attachment was likely characterized by an inflammation response termed the ancestral inflammatory reaction (1, 2). In eutherian pregnancy there is an inflammatory reaction at implantation of the embryo, followed by the establishment of an anti-inflammatory environment necessary to sustain the fetal-maternal interface with the development of the placenta (3). The initial pro-inflammatory environment is necessary for implantation in early pregnancy but leads to termination or premature birth if it is sustained after implantation is complete (4, 5). A key innovation in eutherian pregnancy is the ability to switch off this inflammation after it has been initiated leading to a period of development and growth (1, 6). An anti-inflammatory environment is induced for most of the gestation period in eutherians, followed by a secondary inflammatory phase leading to parturition (7). This long anti-inflammatory period in the middle of pregnancy is characterized by the absence of pro-inflammatory cytokines and an increase in the concentration of anti-inflammatory cytokines such as IL-10 and TGFß (8-10). This extended phase, in which inflammation is suppressed, likely allows for an extended period of placentation, which is a key difference between marsupial and eutherian reproduction (Fig. 1a).

**Fig. 1.**
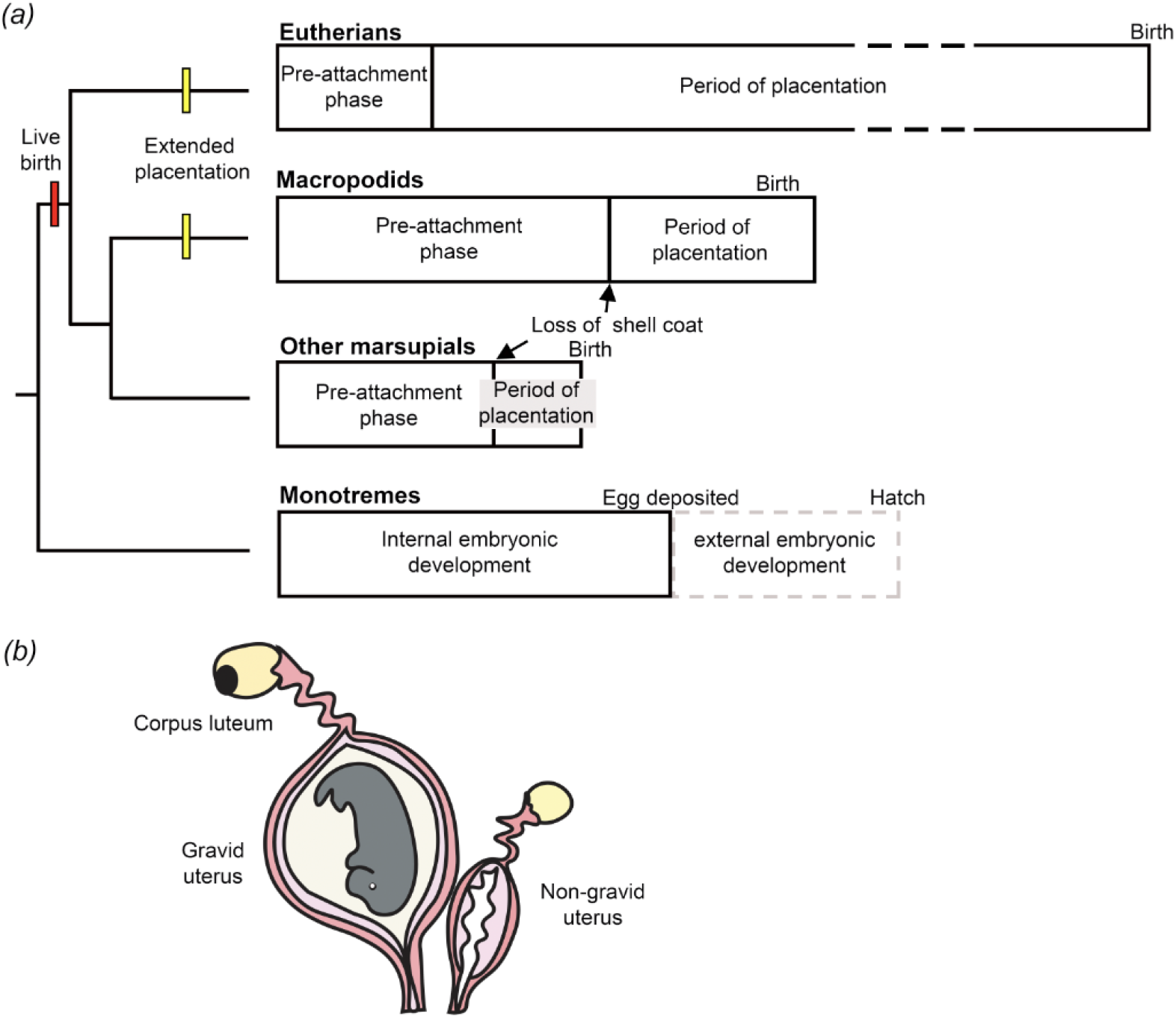
***a)*** Comparison of the length of gestation and proportion of placentation during pregnancy across eutherians, macropodids, other marsupial taxa and monotremes. In general, eutherians have longer gestations than marsupials with gestation extended beyond the length of the oestrous cycle and implantation occurring during early pregnancy. Macropodids have extended placentation compared to other marsupials with gestation almost the same length of time as the oestrous cycle. Most marsupials have shorter gestations than the oestrous cycle, with attachment of the early embryo and placentation occurring in the final 2-3 days of pregnancy. Monotremes have a period of internal embryonic development within the uterus before the eggs are laid into a pouch and external embryonic development continues supported by a complex milk. Yellow bars represent the independent evolution of extended placentation in macropodids and eutherians. The red bar represents the evolution of live birth that occurred once before the split of eutherian and marsupial mammals. ***b)*** The unique reproductive anatomy of macropodids allows us to elucidate the direct effect of the embryo on the uterine environment and gene expression. All marsupials have two separate uteri with an accompanying ovary. The tammar wallaby is monovular with ovulation alternating between the two ovaries. This means that only one uterus becomes gravid each pregnancy (11). A postpartum oestrous and mating normally occurs within an hour of birth. The resulting early blastocyst is then kept in lactationally controlled embryonic diapause. Removal of pouch young leads to reactivation of the embryo from diapause and birth ∼26.5 days later (12). Both uteri are under the same systemic endocrine conditions, but experience different local, unilateral effects based on their proximity to the developing follicle or the corpus luteum (13). Differences between gravid and non-gravid uteri reflect these local effects and the influence of the developing embryo in the gravid uterus (14).

During pregnancy of opossums and other “basal” marsupials (but not macropodids, see below), placentation is brief with shell coat rupture and attachment of the embryo to the uterine epithelium occurring within the last 2-4 days of gestation (Fig. 1a, 15, 16). Phylogenetic evidence supports that opossums represent the more ancestral form of viviparous reproduction (17). Opossums (*Didelphidae*) have retained many of the reproductive features of the first live bearing mammals (17), having a short gestation and short period of fetal-maternal contact, development of a functional yolk sac placenta and no maternal recognition of early pregnancy (17, 18, 19). The grey short-tailed opossum, *Monodelphis domestica*, has a 14-day gestation, with endometrial recognition of pregnancy and an inflammation reaction occurring at attachment on day 12 which continues through the short period of placentation (1,19-21). Opossums lack the anti-inflammatory period after attachment seen in eutherians with an inflammatory cascade leading into parturition (22). The attachment induced inflammation reaction likely limits the length of placentation and therefore pregnancy in marsupials (1). This observation poses the question of whether an anti-inflammatory phase is required for the extension of placentation in mammalian pregnancy in general or whether the moderated inflammation response seen at implantation in eutherian pregnancy is just one method to establish a sustained fetal-maternal interface allowing for extended placentation?

Wallabies offer a unique opportunity to identify how pregnancy and the period of placentation can be extended, in a lineage independent to eutherians. In comparison to other marsupial taxa, macropodids (wallabies and kangaroos) are considered more derived (17, 19). They are the only lineage of marsupials that have an extended period of placentation, greater than the 2-4 days seen in most other marsupial taxa with shell coat rupture occurring 8-10 days before birth in the tammar wallaby (Fig. 1a, 23, 24). The maternal immune profile at attachment and throughout the extended period of placentation in this lineage of marsupials is unknown. The unique reproductive anatomy of macropodids means that only one uterus becomes gravid each pregnancy which allows us to elucidate the direct effect of the embryo on the uterine environment and gene expression (Fig. 1b, 11). To understand the nature of maternal-fetal interactions through extended placentation in the wallaby, we compared gene expression in the uterus of the tammar wallaby from attachment through the late stages of pregnancy. We found that the molecular environment at the maternal-fetal interface during maternal-fetal attachment in the tammar wallaby is homologous to that which occurs at attachment in the opossum and implantation in eutherian pregnancy. However, this pro-inflammatory state at attachment is followed by a uniquely moderated inflammatory profile with some key mediators of inflammation not expressed at all throughout tammar wallaby pregnancy. We argue that it is this moderation of an inflammation reaction at attachment and the subsequent suppression of inflammation that has facilitated extension of placentation in the macropodid lineage.

## Results

### The tammar wallaby uterus undergoes vast remodeling during attachment and placentation

Histology of the endometrium shows that some changes to the vasculature and gland structure are driven by maternal cycling due to similar changes occurring in both gravid and non-gravid uteri at key timepoints of pregnancy (Fig. 2). Before attachment the endometrium from both non-gravid and gravid uteri has columnar epithelium with basally located nuclei. Capillaries are present underlying the luminal epithelium. After attachment there is substantial glandular and vascular development in both gravid and non-gravid uteri. The glandular epithelium is more cuboidal in structure with dilated gland openings and pink eosinophilic staining substances within the lumen. At this pre-attachment stage there are similarities in morphology of gravid and non-gravid uteri from the same timepoints of pregnancy. This supports the suggestion that some changes to the endometrium occur due to the systemic hormonal changes that occur during pregnancy rather than the presence of the fetus (25, 26, 27).

**Fig. 2.**
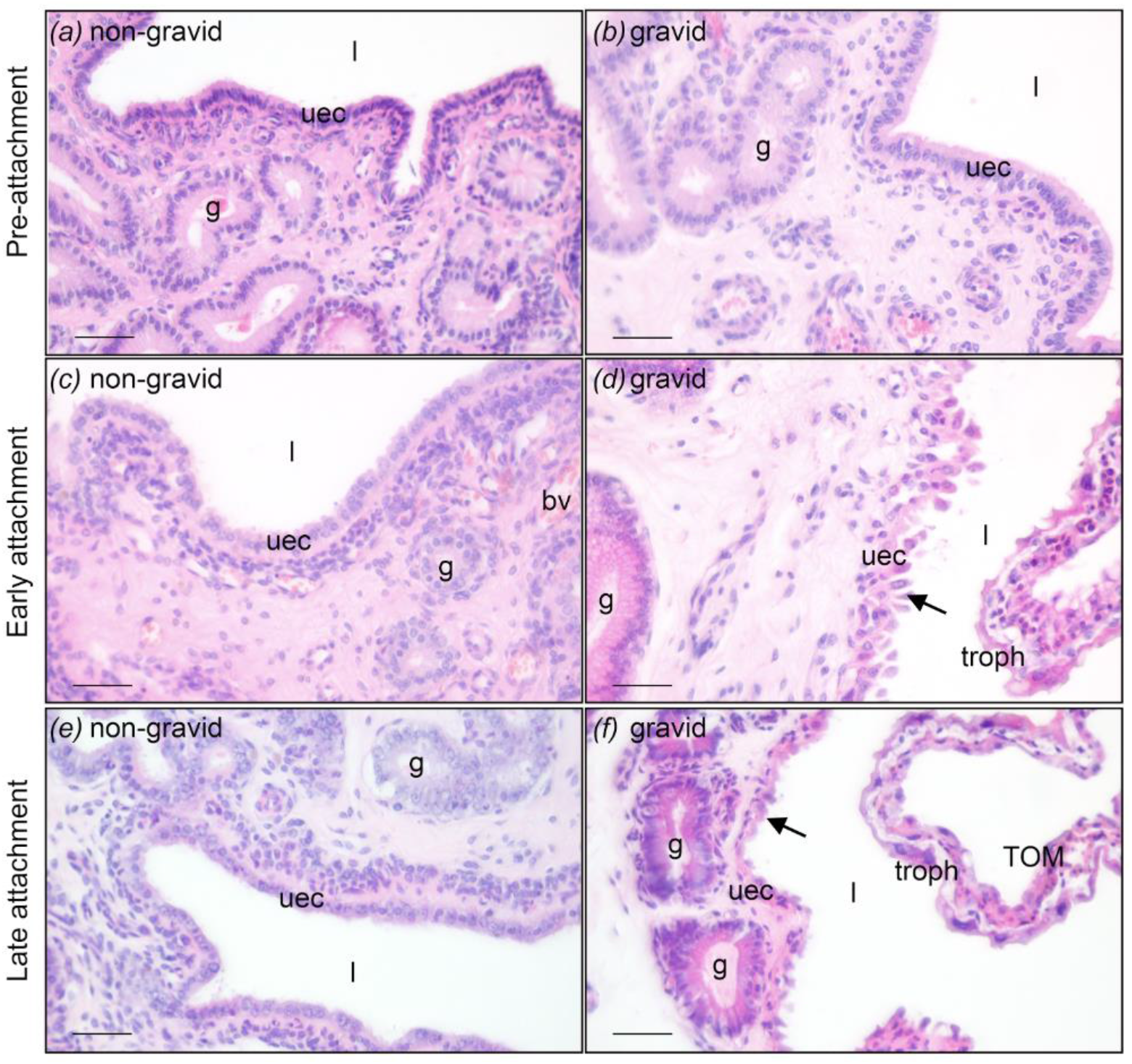
Histological comparison of non-gravid and gravid uterus throughout pregnancy using hematoxylin and eosin staining. ***a)*** Uterine sections from non-gravid uteri and ***b)*** gravid uteri at day 14: pre-attachment of the blastocyst and loss of the shell coat (days 14-16, n=2) show similar morphology of uterine epithelium. ***c)*** and ***d)*** Early attachment, after loss of the shell coat and formation of the placenta (days 20-23, n=3) the uterine epithelium of gravid uteri has rounded apices and appears to be blebbing (arrow). ***e)*** and ***f)*** Late attachment, immediately before birth (days 25-26, n=2) the uterine epithelium is short and rounded (arrow) in gravid uteri compared to non-gravid.These morphological changes to the uterus through attachment in the tammar wallaby are similar to the short period of attachment in the opossum, and implantation in eutherians. Staging is listed as days post copulation. The uterus is lined by luminal epithelium (uec) and the underlying stroma consists of connective tissue with uterine glands (g) embedded throughout. bv: blood vessel, l: uterine lumen, uec: uterine epithelial cells, g: gland, troph: trophoblast, TOM: trilaminar portion of the yolk sac. All scale bars are 50μm.

After attachment the uterine environment is differentially re-modelled in the presence of the fetus. After attachment the endometrium is greatly folded with regions closely interdigitated with fetal membranes (Fig. 2d). The luminal epithelium changes its structure, flattening out with rounded apices. This rounded structure is a feature of cellular blebbing which can be caused by detachment of adhesive structures along cell membranes (28). The domed shape and blebbing of the uterine epithelium is present throughout the entire period of attachment also seen in the final days of pregnancy (Fig. 2f). However, it is not seen in non-gravid uteri (Fig. 2e) supporting the conclusion that it is the presence of the developing fetus and the placenta that influences this maternal recognition of pregnancy. The extent of blood vessel and glandular development is increased in gravid uteri compared to non-gravid uteri at the same stage (13), suggesting that these changes facilitate the formation of the placenta.

### Reproductive state and timepoint of pregnancy have substantial impacts on uterine gene expression

Transcriptomic data shows that the stage of pregnancy (pre-attachment, post-attachment, end of pregnancy) has the biggest impact on differences in uterine gene expression followed by the reproductive state (gravid and non-gravid uteri) (Fig. 3a). Pairwise comparisons show differentially expressed genes between non-gravid and gravid uteri at each timepoint of pregnancy (Fig. 3b). Samples from the pre-attachment stage of pregnancy (day 14 gravid and day 14 non-gravid) distinctly separate from the post-attachment stages of pregnancy and post-partum uteri (Fig. 3a) with hundreds of differentially expressed genes in pairwise comparisons (Fig.3c). There is distinct separation of samples from pre-attachment (day 14) and early attachment (day 20) gravid and non-gravid uteri. The greatest separation is seen between gravid and non-gravid uteri from late attachment (day 25). This suggests that the presence of the fetus has more influence on gene expression differences during the period of attachment and before birth than the pre-attachment stages of pregnancy. Non-gravid samples from the latest stage of pregnancy (day 25 non-gravid) cluster close to samples from pre-ovulation uteri (PO; Fig. 3a) and do not have any significantly differentially expressed genes in pairwise comparisons (Fig. 3d). Pre-ovulation uteri samples are from the uterus contralateral to the post-partum uterus within 24 hours of birth. Ovulation occurs from the ovary contralateral to the post-partum uterus within 40 hours of birth and the embryo enters the uterus ipsilateral to this ovary up to 2 days later.

**Fig. 3.**
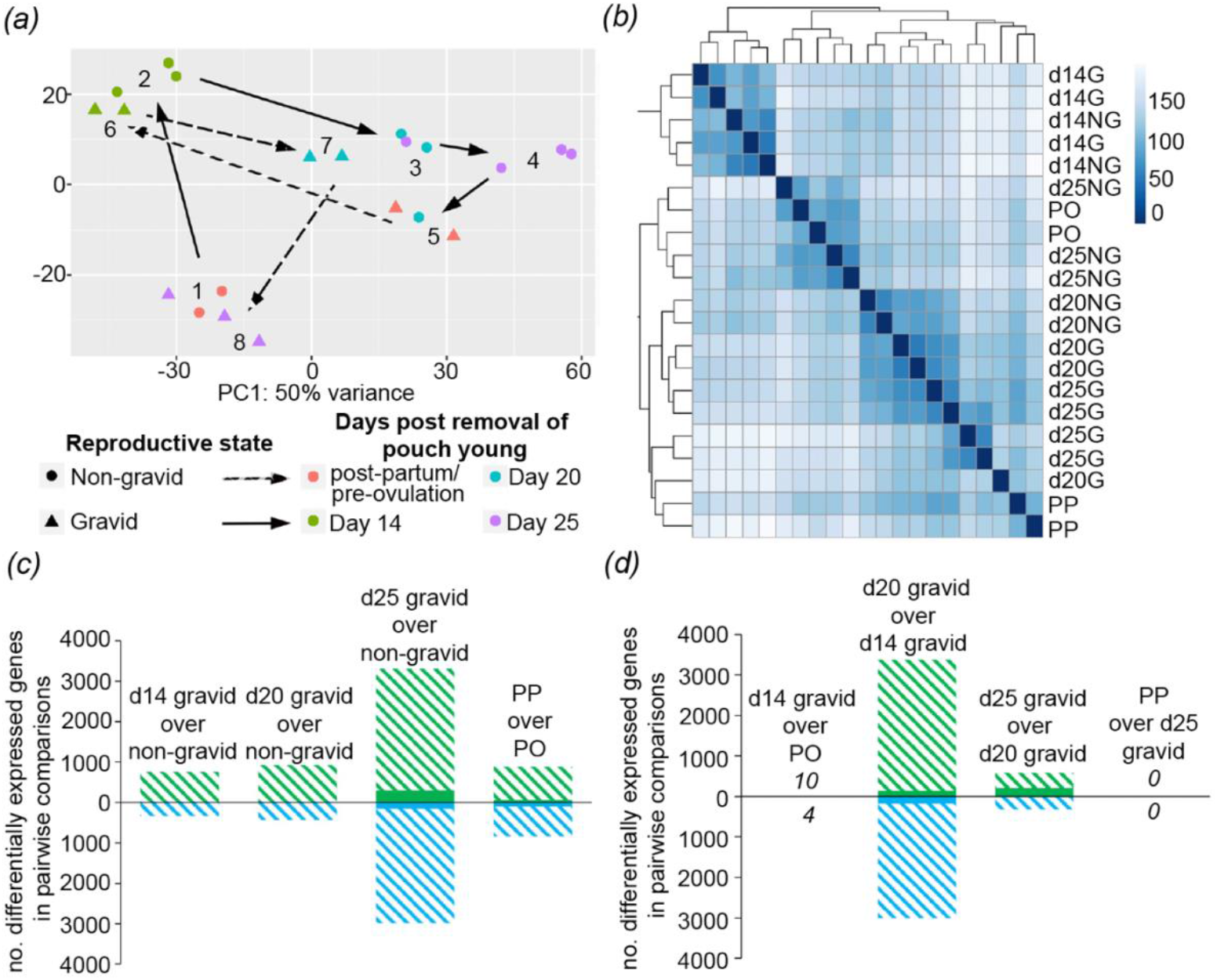
Correlations of transcriptome samples. ***a)*** Principal component analysis (PCA) of uterine transcriptomes of different reproductive states and timepoint during pregnancy. ***b)*** Heatmap showing pairwise comparisons. ***c)*** Number of differentially expressed genes in pairwise comparisons of gravid over non-gravid uteri. Bars show the number of significantly upregulated (hash) and downregulated (blue) genes in gravid compared to non-gravid uteri from pre-attachment (d14), early attachment (d20), late attachment (d25), pre-ovulation uterus (PO) and post-partum uterus (PP). Hashed bars represent the total number of significantly differentially expressed genes, while the solid bars represent significantly differentially expressed genes with a fourfold difference in expression. ***d)*** Number of differentially expressed genes in pairwise comparisons of pre-ovulation (PO), pre-attachment (d14) post-attachment (d20), late attachment (d25) and post-partum (PP) uteri. Hashed bars represent the total number of significantly differentially expressed genes, while the solid bars represent significantly differentially expressed genes with a fourfold difference in expression.

There are thousands of differentially regulated genes after the shell coat is lost, and attachment occurs (comparing pre-attachment to early attachment). With loss of the shell coat and attachment, the endometrium comes into direct contact with the choriovitelline placenta. This process causes endometrial remodeling and likely complex signaling/ communication between mother and fetus. Gene Ontology (GO) analysis shows that at early attachment compared to pre-attachment there is a significant enrichment of genes associated with cell adhesion and extracellular matrix organization with clear clustering of GO terms based on biological processes (Fig. 4a). This aligns with the reinforcement of cell adhesion and remodeling of the plasma membrane that occurs at embryo attachment with the development of an epitheliochorial placenta in this species (29, 30). There is a cluster of overrepresented gene ontology terms related to the regulation of cell migration which is interesting as there is no migration of invasion of uterine epithelial cells with non-invasive placentation in the tammar wallaby. It is more likely that these gene are associated with changes to the endometrial stroma. We found an upregulation of genes associated with tight junction and gap junction proteins (TJP1, CLDN7, CLDN23, GJA5) and cadherin associated proteins (CTNNA1, CDH3, CDH7) which play a role in maintaining the adhesion of uterine epithelium (reviewed in 31). There is also an upregulation of genes associated with aquaporin (AQP1) and mucins (MUC1, MUC7), which play a role in maintaining polarization of uterine epithelium and movement of proteins during implantation in species with invasive placentation (32, 33). There is an upregulation of proteases expressed in the endometrium such as matrix metallopeptidases (MMP25, MMP7, MMP15) which may play a role in the breakdown of the shell coat that occurs at attachment in the tammar wallaby or dissolving of the endometrium at this stage of pregnancy. There is also enrichment of genes associated with angiogenesis and regulation of vascular development (VEGF, ESM1) which supports the substantial increase in vasculature that occurs throughout the endometrium during the later stages of pregnancy.

**Fig. 4.**
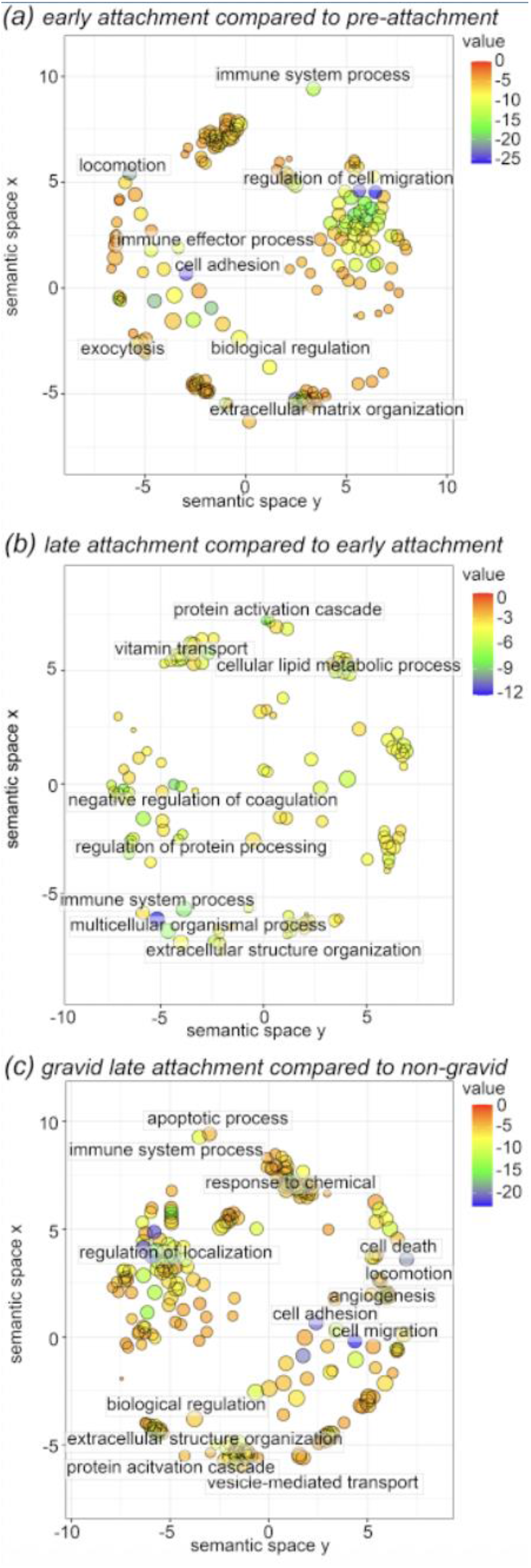
GOrilla gene ontology (GO) enrichment analysis of differentially expressed genes using scatterplots from REVIGO for biological processes. ***a)*** significantly over-represented GO terms from the list of significantly upregulated genes at early attachment (d20) compared to pre-attachment (d14). ***b)*** significantly over-represented GO terms from the list of significantly upregulated genes at late attachment (d25) compared to early attachment (d20). ***c)*** significantly over-represented GO terms from the list of significantly upregulated genes in gravid uteri at late attachment (d25) compared to non-gravid uteri from the same stage.

The late attachment timepoint of pregnancy, 1.5-2 days before birth (day 25) shows the greatest number of differentially expressed genes correlating to the extensive morphological and physiological changes that occur to the endometrium at this timepoint. The most over-represented GO terms (top 10 gene ontology terms ordered by p-value) upregulated when comparing the early and late attachment timepoints which span the length of placentation, are associated with fibrinolysis, regulation of coagulation and wound healing (Fig. 4b). Clustering of gene ontology terms is not as clear when comparing these timepoints suggesting that the time of placentation features many of the same changes to biological processes. At late attachment, the uterus is preparing for parturition, involving further substantial reorganization of the endometrium and detachment of the fetus at birth. Although placentation is non-invasive in macropodids, there is vast vascular remodeling throughout the period of placentation (34) and tissue injury at the time of birth as suggested by the upregulation of genes associated with wound healing. Although there are a substantial number of differentially expressed genes throughout the period of attachment (upregulated on day 25 gravid compared to day 20) there are no differences in gene expression between uteri from the final days of pregnancy (day 25) and post-partum (PP) suggesting that it takes longer than 24 hours for the endometrium to transition back to a non-gravid state. The pre-ovulation uteri (PO) have very few differentially expressed genes and no enriched GO terms when compared to gravid pre-attachment uteri (d14). This suggests that there are few gene expression differences in early pregnancy before attachment while the developing fetus is surrounded by a shell coat. These early stages are characterized by minor changes to the morphology of the endometrium and no direct contact between the maternal endometrium and developing fetus which could explain the low number of differentially expressed genes.

### Marsupial attachment is homologous to eutherian implantation

We investigated the expression of key implantation markers from eutherian pregnancy in the tammar wallaby. Expression of Heparin-binding EGF-like growth factor (HBEGF) and Mucin 1 (MUC1) are upregulated after the loss of the shell coat and maintained throughout the period of attachment (d20 and d25; Fig. 5). MUC1 expression increases further in post-partum uteri but HBEGF expression decreases after birth. MMP7 expression is upregulated in tammar wallaby endometrium during the period of attachment. Immunohistochemistry shows that MMP7 staining is localized almost exclusively in the nucleus of the uterine epithelium which suggests that it has a gene regulatory function during late attachment (d25; Fig. 5f). In the fetal membranes staining is both nuclear as well as cytoplasmatic. There is no staining in the uterine endometrium from the pre-attachment stage (d14) of pregnancy which correlates with the low expression levels at this reproductive stage (Fig. 5e).

**Fig. 5.**
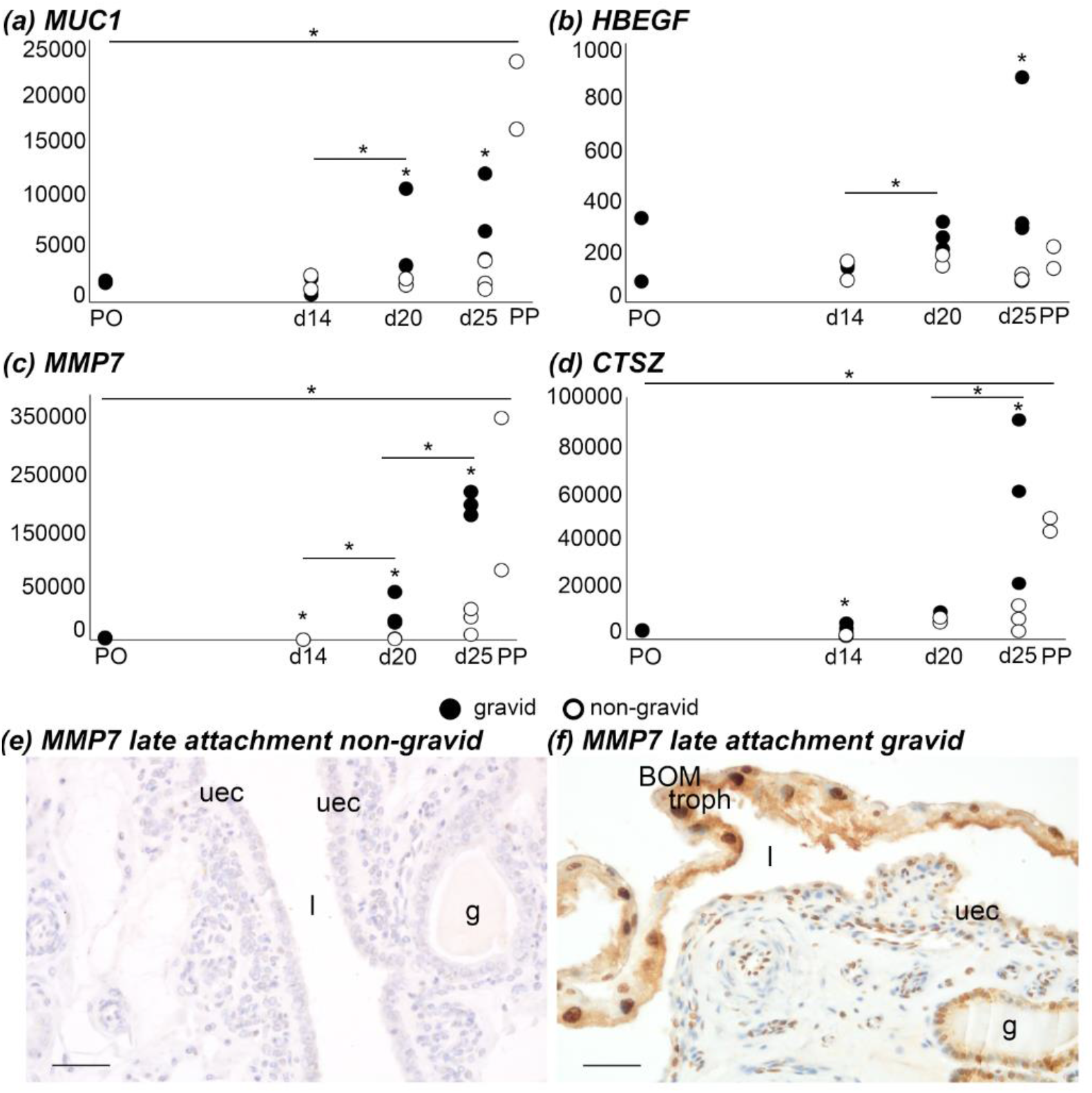
Transcriptomic gene expression of implantation markers with day of pregnancy on the x-axis and relative read counts on the y-axis. ***a)*** MUC1, ***b)*** HBEGF and proteases ***c)*** MMP7 and ***d)*** Cathepsin Z in gravid and non-gravid uteri at different timepoints during tammar wallaby pregnancy. Uteri from pre-ovulation (PO), pre-attachment (d14), post attachment (d20), late attachment (d25) and post-partum (PP). Asterisks show significance (p<0.05). ***e)*** and ***f)*** There was no immunolocalization of the protease MMP7 in non-gravid uteri compared to that in the uterine epithelium and fetal membranes in gravid uteri at late attachment (d25).Nuclei are blue due to hematoxylin counter staining and the immunostaining signal Is brown due to 3,30-diaminobenzidine (DAB). l: uterine lumen, uec: uterine epithelial cells, g: gland, troph: trophoblast, BOM: bilaminar portion of the yolk sac. All scale bars are 50μm.

There is a controlled upregulation of expression of cathepsins in tammar wallaby endometrium during the period of attachment (CTSZ, CTSC, CTSD, CTSB) along with Cystatin C (CST3). This expression pattern is similar to that which occurs in eutherian implantation (mice, sheep and pig) where cathepsins are likely involved in remodelling of the endometrium through degrading the extracellular matrix (35-37).

### There is a modified inflammatory gene expression profile as pregnancy progresses

To determine the inflammatory gene profile during pregnancy and how it is affected by the presence of a fetus we focused on GO terms associated with an inflammatory response between non-gravid samples and gravid samples at matching timepoints. There was no enrichment of GO terms related to an inflammatory response between gravid and non-gravid samples before attachment (d14) suggesting that the presence of a shell coat around the developing fetus prevents a maternal inflammatory response in early pregnancy.

There is a significant overrepresentation of GO terms related to an immune response before birth (d25) in the gravid uterus compared to the non-gravid uterus at the same timepoint (Fig. 4c). This provides evidence for the influence of the fetus on endometrial gene expression after attachment inducing a moderated inflammatory response with some inflammatory genes being upregulated throughout the period of placentation whilst others are suppressed or show a modified expression. There is an over-representation of genes associated with an inflammatory response and immune system processes such as LIF - leukemia inhibitory factor, complement genes (C2, C3, C9), Major histocompatibility complex, class i, a, (HLA-A), Interleukins and their receptors (IL6, IL1RAP, IL1R1, IL12A) and genes associated with prostaglandin synthesis (PTGS2, PTGES, PTGDR, PTGFR). There is also an increase in the expression of genes involved in neutrophil activation and neutrophil mediated immunity such as proteases (PRSS2) and cystatins (CSTB, CST3).

To further test the hypothesis that an inflammatory reaction occurs at attachment in the tammar wallaby and is moderated throughout the period of placentation we assessed the expression of marker genes of inflammation. We found that a variety of inflammatory genes are upregulated at key time points of pregnancy. Cytokines, such as IL6, are upregulated before attachment (d14) and birth (d25) and maintained in post-partum uteri (PP; Fig. 6a). LIF expression is upregulated after attachment (d20) and maintained during the period of placentation (d20 and d25; Fig. 6b). LIF expression is also maintained in both pre-ovulation (PO) uteri and postpartum uteri (PP). The upregulation of IL6 and LIF at attachment and before birth suggests that an inflammatory response occurs at these timepoints.

**Fig. 6.**
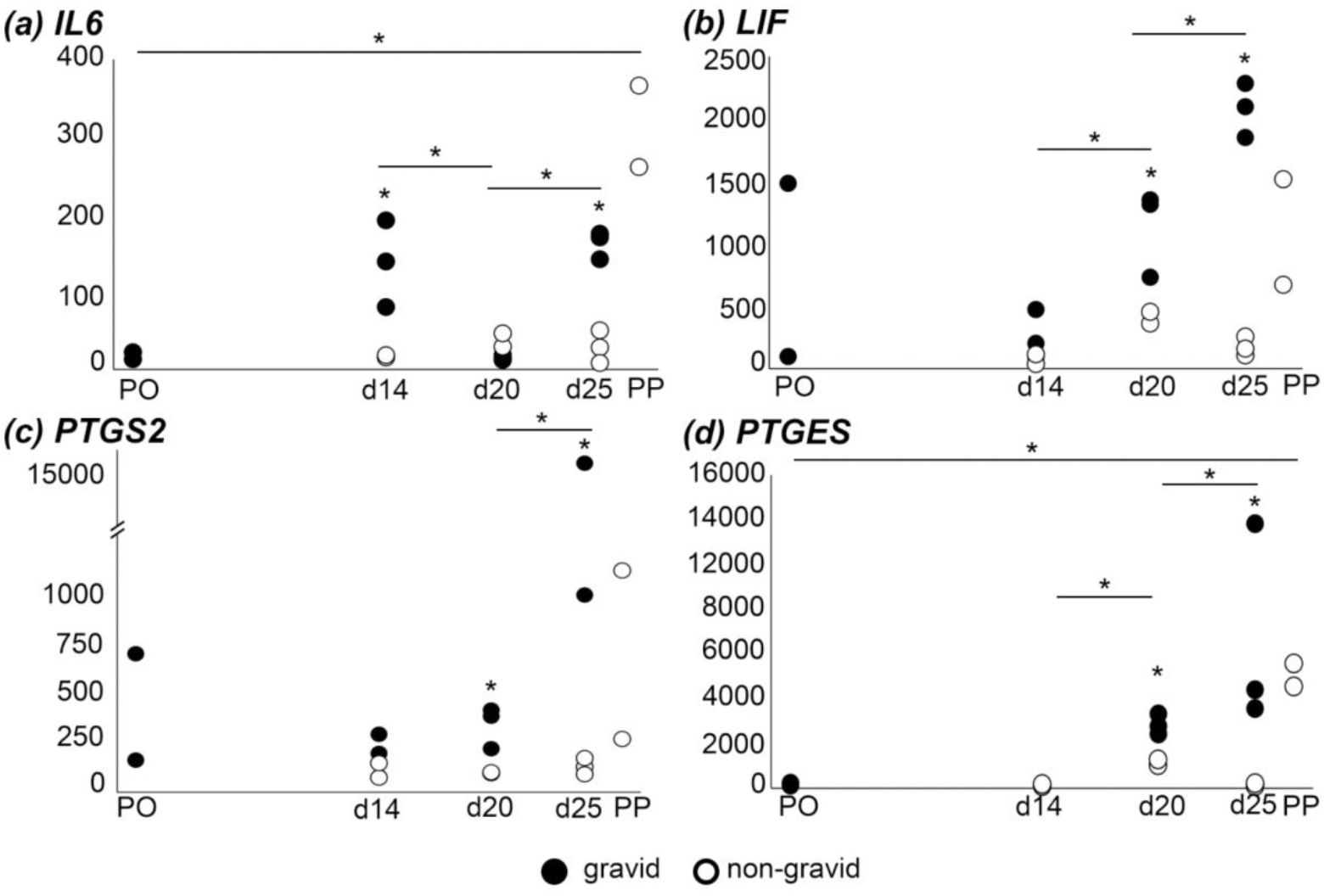
Transcriptomic gene expression of mediators of inflammation with day of pregnancy on the x-axis and relative read counts on the y-axis. ***a)*** IL6, ***b)*** LIF, ***c)*** PTGS2 ***d)*** PTGES in gravid and non-gravid uteri at different timepoints during tammar wallaby pregnancy. Uterine samples were collected from the pre-ovulation (PO), pre-attachment (d14), early attachment (d20), late attachment (d25) and post-partum (PP) stage of pregnancy. Asterisks show significance (p<0.05).

There is an enrichment of genes associated with an immune effector process at early attachment (d20) compared to pre-attachment (d14; fig. 4a). During early attachment there is an enrichment of genes associated with an inflammatory response with an upregulation of acute phase proteins known to be involved in an inflammatory response such as Coagulation factor viii (F8) and interleukins (IL34). There is an upregulation of genes associated with fibrinogen (FGG, FGA and FGB) at pre-attachment (d14), followed by suppression during early attachment (d20) and further upregulation at late attachment before birth (d25). This suggests that there is a moderated inflammatory reaction through the period of attachment as the fetus grows and the fetal maternal interface is developed before birth.

In the tammar wallaby, genes associated with prostaglandin synthesis are upregulated leading up to birth which is known to facilitate parturition. PTGS2 and PTGES expression in gravid uteri is upregulated at late attachment, before birth (d25; Fig. 6c, d) as prostaglandins are in the yolk sac and yolk sac fluid and circulation (38, 39). PTGS2 is upregulated before birth then reduces in post-partum uteri (PP). Circulating prostaglandins drop dramatically within an hour of birth (39). This gene expression pattern aligns with the spike in concentration of PGF2α, PGE2 and 6-keto-PGF1 seen at birth in tammar wallaby uterine and fetal tissues (38, 39). Circulating prostaglandins then drop dramatically within an hour of birth (39).

The differential expression of some inflammatory genes and not others throughout pregnancy suggests that a moderated inflammatory response occurs in the tammar wallaby. Despite the upregulation of known inflammatory markers, other key regulators of an inflammatory response such as IL17A, IL10, IL1A show very low raw reads with no differential gene expression through pregnancy. The low expression of these key inflammatory genes suggests that only a partial inflammatory response occurs during attachment in the tammar wallaby.

## Discussion

We show that during tammar wallaby pregnancy there is a moderated inflammatory response with some inflammatory genes expressed at key time points of gestation. Pro-inflammatory cytokines including *IL6*, are expressed before embryo attachment, *IL12A* and *LIF* throughout the period of placentation and prostaglandins before birth. These results are consistent with specific inflammatory genes being co-opted into regulation of tammar pregnancy, while a generalized inflammatory response is being suppressed or modified. We argue that modulation of inflammation is a key innovation that allows for the extension of placentation in mammals. This progressive inflammation reaction differs to the ancestral attachment reaction seen in opossum pregnancy and the implantation inflammatory reaction in eutherian pregnancy (Fig.7).

### The period of attachment is homologous to eutherian implantation

Several morphological changes to the uterus through attachment in the tammar wallaby suggests that this period of close maternal-fetal apposition is homologous to the short period of attachment in the opossum, and implantation in eutherians.

Remodeling of the endometrium and gene expression patterns during the period of attachment and placentation mirror the changes seen during attachment in the opossum (1, 41, 42) and implantation in eutherian mammals (43, 44). Key implantation markers, Heparin-binding EGF-like growth factor (HBEGF) and Mucin 1 (MUC1) are upregulated after the loss of the shell coat and maintained throughout the period of placentation in the tammar wallaby. Similarly, in the opossum, there is a moderate increase in expression of MUC1 following attachment and a further increase in expression through the final days of pregnancy (1). This increased expression pattern is also seen in eutherian species such as rodents and humans where MUC1 is upregulated in uterine epithelium during the window of implantation. However, its expression is lost in maternal epithelium close to the points of embryo attachment (45, 46). During eutherian implantation an inflammatory gradient created by cytokines and chemokines facilitates the formation of a mucin layer and remodeling of the uterine epithelium which is necessary for apposition and adhesion of the blastocyst at implantation (reviewed in 3). Despite the modification of the inflammation response seen during pregnancy in the tammar wallaby, the differential expression of key implantation markers throughout the extended period of placentation in the tammar wallaby provides evidence for the period of attachment being homologous to the window of implantation in eutherian pregnancy.

The process of attachment and implantation of the embryo is associated with remodeling of the endometrium. Many features of inflammation are shared with tissue remodeling and so may play a beneficial role in the process of attachment and implantation (47). For example, during pregnancy, mast cells are activated in response to sex steroids (estradiol and progesterone), to release mediators such as histamine, VEGF, proteases, and metalloproteinases (MMPs). These mediator proteins and compounds contribute to an inflammation response (48-51). MMPs also degrade the extracellular matrix of the endometrium and are important regulators of vascular and uterine remodeling (reviewed in 48, 52). High levels of MMP2 protein have been identified in uterine flushings from tammar wallabies in late pregnancy (53). There is an upregulation of matrix metallopeptidases (MMP25, MMP7, MMP15) in tammar wallaby endometrium throughout placentation. The maternal expression of MMP7 during tammar wallaby pregnancy with localization of Immunohistochemistry (IHC) staining to nuclei in the endometrium suggests it has a gene regulatory function there. However, there is also strong staining in the cytoplasm of the trophoblast which suggests a different secretory function for embryonic tissues. Key markers of implantation such as mucins and metalloproteinases, show a moderated expression throughout the period of placentation in the tammar wallaby with similarities to eutherian species highlighting that this period is homologous to the period of implantation in eutherian pregnancy.

### *Only the ghost of an inflammation response persists in* tammar *wallaby pregnancy*

Opossums experience an acute inflammatory reaction at attachment which likely limits the extension of placentation beyond a few days. This is characterized by a sustained upregulation of genes involved in an inflammatory reaction in the uterus during placentation such a proinflammatory cytokines (IL6, TNF, IL17A) and prostaglandins (PTGES and PTGS2; 1). The pro-inflammatory cytokines, IL17A and IL6 are highly expressed by the trophoblast giant cells at embryo attachment (40).Eutherian implantation has the relatively full gamut of inflammatory signaling expected of a mucosal tissue but is missing IL17A signaling. The suppression of IL17A signaling is achieved by decidual stromal cells, and results in downstream prevention of neutrophil recruitment to the endometrium, which is likely important for establishing placentation in eutherians (2, 40).

The inflammatory environment at implantation across eutherian lineages is mostly conserved (Fig. 7). Inflammatory cytokines LIF, IL6, IL1A and IL1B and prostaglandins are expressed at the fetal-maternal interface at implantation across eutherian lineages including the more basally branching eutherian clades such as Xenarthra (which includes the armadillo; 2, 40, 54, 55). LIF, which is absent during placentation in the opossum, plays an important role in decidualization of endometrial stomal cells during eutherian implantation (56). Before implantation in eutherians there is an upregulation of VEGF which is a potent inducer of increased vascular permeability associated with angiogenesis and inflammation (57, 58). This is followed by an estrogen induced increase in blood flow resulting in uterine oedema which creates an optimal environment for endometrial remodeling and implantation (59). In eutherian pregnancy, the inflammatory environment preceding birth also promotes contraction of the uterus (6). Prostaglandins are key markers of acute inflammation playing a role in increasing vascular permeability (60). Some of them, such as PGF1a, also induce contraction of the myometrium contributing to the process of parturition in eutherian and marsupial species (61, 62, reviewed in 63).These features of an inflammation response may have been co-opted into important roles of pregnancy such as increased vascular permeability, endometrial remodeling, and initiation of birth.

**Fig. 7.**
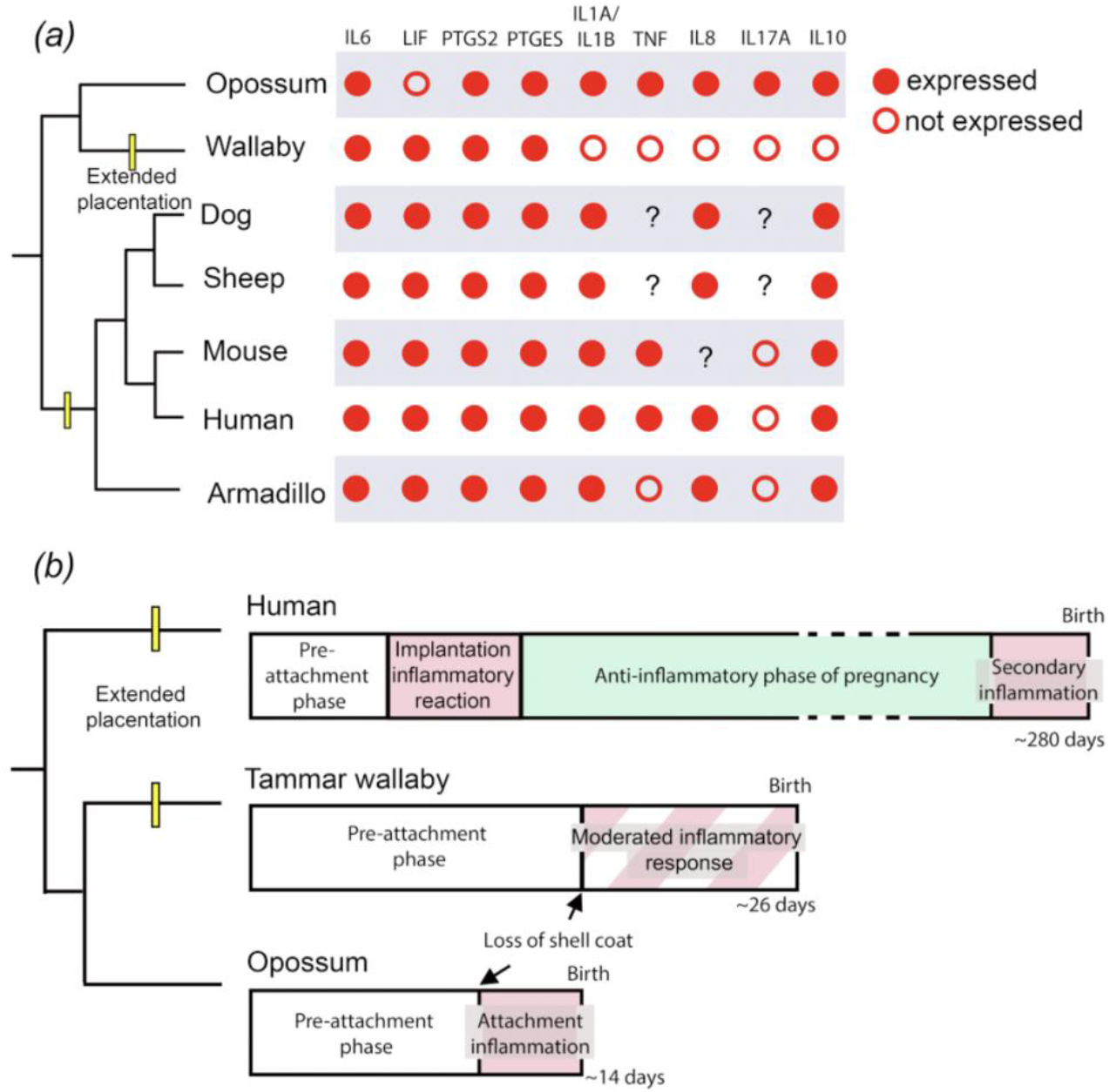
Comparison of the inflammatory markers and the inflammation profile during pregnancy across Theria. ***a)*** Comparison of key mediators of inflammation at attachment across Theria (reviewed in 2, 40). Pro-inflammatory cytokines such as IL17A, 1L1A, IL1B, IL8, IL10 and TNF are expressed at low levels in tammar wallaby endometrium but show no differential expression throughout pregnancy. ***b)*** Comparison of the length and maternal immune profile across eutherian and marsupial pregnancy showing the difference in immune profile across human, tammar wallaby and opossum pregnancy.Eutherian pregnancy is characterized by two pro-inflammatory phases at implantation and before birth. Macropodids such as wallabies have extended placentation compared to other marsupials with a moderated inflammation response occurring during the period of placentation. Adapted from (1).Yellow lines represent the independent evolution of extended placentation in macropodids and eutherians.

In contrast to eutherians, we found that some inflammatory genes are expressed at attachment and throughout placentation in tammar wallaby pregnancy but not others. Genes that are expressed in the ancestral inflammatory attachment reaction such as IL17A, IL1A, and IL1B are not expressed during pregnancy in the tammar wallaby.The absence of key mediators of inflammation suggests that there is no complete inflammation reaction at attachment in the tammar wallaby. The loss of expression of particular inflammatory genes through the period of placentation likely allows for the extension of placentation in macropodids. In comparison to both the opossum and eutherian state, many of the downstream inflammatory genes are not differentially expressed at attachment in tammar wallaby pregnancy (Fig. 7). The genes that are differentially expressed are regulators of an inflammatory response including Interleukin-6 family cytokines. The genes that have been retained during tammar wallaby pregnancy are specific to inflammation and regulate inflammatory processes such as increased vascular permeability, remodeling of the extracellular matrix, oedema, and tissue repair. This provides support for the hypothesis that some features might be beneficial and being integrated into the physiology of implantation and attachment during pregnancy. This moderated expression of key mediators of inflammation is unique to macropodid pregnancy, likely playing a role in extending the period of placentation in this marsupial lineage.

### Building a general model for the evolution of extended placentation

Our findings suggest that the moderation of inflammation at embryo attachment is a general requirement to extend the period of placentation in mammals. The short period of attachment experienced by the first live bearing mammals was likely characterized by an inflammation response similar to what occurs during opossum pregnancy (1, 2). This inflammatory reaction at attachment limited the length of placentation due to a sustained influx of pro-inflammatory factors contributing to the onset of parturition after only a few days of maternal-fetal interaction.

During eutherian pregnancy a short pro-inflammatory environment is necessary for embryo implantation but is switched off after implantation for most of placentation (4, 6). In macropodids, another mammalian lineage that has extended placentation, we show that they also moderate inflammation, by silencing key inflammatory genes throughout embryo attachment and the period of maternal-fetal contact. We propose that moderation of inflammation after embryo attachment is key to extending mammalian pregnancy during the period of maternal-fetal interaction.

## Methods

### Sample collection and design

Tammar wallaby uterine tissue was collected from our breeding colony at the University of Melbourne. All experiments were approved by the University of Melbourne Animal Ethics Committees and followed the National Health and Medical Research Council (2013) guidelines. The endometrium was separated from the myometrium and fetal tissues by dissection and samples were stored for RNA analysis, histology, and western blot analysis.

### Histology and immunostaining

Samples of gravid and non-gravid uteri were collected for histological analysis at pre-attachment (days 14-16, n=2), early attachment (days 20-23, n=3), late attachment (days 25-26, n=2) and post-partum (n=2) stages of pregnancy based on the number of days after the removal of pouch young. Fixed tissues were embedded in paraffin, sectioned, deparaffinized, cleared and then stained with hematoxylin and eosin following a standard protocol (64).

The expression of MMP7 was localized using immunohistochemistry. Uterine samples were fixed, embedded in paraffin, sectioned, de-paraffinized, cleared, and then incubated in primary antibody specific to the protein of interest (rabbit anti-MMP7 antibody #NBP1-99123 (1.28 mg/mL; Novus Biologicals). Sections were washed and incubated with a secondary antibody (Polymer-HRP Anti-rabbit and DAB following kit protocol (DC EnVision + System, HRP; DAKO). Slides were counterstained with hematoxylin, dehydrated and cover slipped for imaging using an Olympus BX53 with DP74 camera and cellSens Imaging Software (Olympus). The specificity of the primary antibody was confirmed with Western blot analysis (Appendix methods and results).

### RNA sequencing and analysis

For RNA-seq analysis we examined uterine tissue from non-gravid and gravid uteri from pregnant females based on the number of days after the removal of pouch young. The stages included pre-ovulation (sample collected from the uterus contralateral to the post-partum uterus within 24 hours of birth; n=2), pre-attachment (day 14; gravid n=3, non-gravid n=2), early attachment (day 20; gravid n=3, non-gravid n=2), late attachment (day 25; gravid n=3, non-gravid n=3) and post-partum (sample collected within 24 hours of birth; n=2). Illumina sequencing libraries were generated by the Ramaciotti Center for Genomics (UNSW Sydney, Australia) using the TruSeq Stranded mRNA prep and sequenced with NovaSeq S1 100bp sequencing. Raw reads were aligned to the tammar wallaby genome v3.0 using hisat2 and reads were counted using htseq-count. Hierarchical clustering and principal component analysis (PCA) was performed in the R *stats* package (65) to confirm that there were general patterns in gene expression between treatment groups. We compared differential gene expression analysis between non-gravid PP, d14, d20, d25 and pre-ovulation (PO), gravid d14, d20, d25 uteri using DeSeq2 default parameters (66, Appendix Table S1-S8). Gene ontology was analyzed using GOrilla and visualized with REVIGO (Appendix Table S9-S21, 67).

## Supporting information

Appendix methods and results

## Acknowledgments

This research was funded by an Australian Research Council Discovery Project (DP200100344) to O.W.G., M.B.R and G.P.W and a John Templeton Foundation Grant no 54860 to G.P.W.

## Author contributions

O.W.G, M.B.R and G.P.W designed the research, O.W.G, M.B.R collected samples, O.W.G and J.S.D analyzed the data, J.S.D, O.W.G, M.B.R and G.P.W wrote the paper. All authors declare no competing interests. This paper was a direct submission to PNAS.

## Notes

### Competing Interest Statement

The authors have declared no competing interest.

