## Appendix methods and results for "The extension of mammalian pregnancy required taming inflammation; independent evolution of extended placentation in the tammar wallaby"

### Supplementary material

#### Supplementary methods

*Sample collection and design*
Tammar wallaby uterine tissue was collected from our breeding colony at the University of Melbourne. All experiments were approved by the University of Melbourne Animal Ethics Committees and followed the National Health and Medical Research Council (2013) guidelines. The endometrium was separated from the myometrium and fetal tissues by dissection and samples were stored for RNA analysis, histology, and western blot analysis.

*Histology and immunostaining*
Samples of gravid and non-gravid uteri were collected at pre-attachment (days 14-16, n=2), early attachment (days 20-23, n=3), late attachment (days 25-26, n=2) and post-partum (n=2) stages of pregnancy based on the number of days after the removal of pouch young. Tissue was fixed in paraformaldehyde and dehydrated through a graded ethanol series, cleared in toluene, and embedded in paraffin. Cross sections of 7 µm were cut using a Leica microtome and mounted on polysine slides (Thermo Scientific). Hematoxylin and eosin staining was completed following a standard protocol (64).

The expression of MMP7 was localized using immunohistochemistry. Uterine samples were sectioned as above from gravid and non-gravid uteri at pre-attachment (days 14-16, n=2), early attachment (days 20-23, n=3), late attachment (days 25-26, n=3). Slides were de-paraffinized in three washes of xylene (3 min) then 100% ethanol (3 min). Antigen retrieval was performed in citrate buffer (12 mM sodium citrate, pH 6.0, 98 °C, 1 h). Slides were incubated in primary antibody overnight at 4°C (rabbit anti-MMP7 antibody #NBP1-99123 (1.28 mg/mL) Novus Biologicals. Each slide was washed with PBS for 10 min and incubated with Polymer-HRP Anti-rabbit for 1 hr and DAB for 5 min following kit protocol (DC EnVision + System, HRP; DAKO).  Slides were counterstained with hematoxylin, dehydrated and cover slipped for imaging using an Olympus BX53 with DP74 camera and cellSens Imaging Software (Olympus).

*Western blotting*
Sections of uterine tissue for western blot analysis were snap frozen and stored at -80°C. Samples from both gravid and non-gravid uteri at each stage of pregnancy (n=1) were homogenized by grinding tissue in a mortar with liquid nitrogen. Tissue was added to 1 X RIPA buffer containing protease inhibitor (RIPA Lysis Buffer System; Santa Cruz Biotechnology). Protein samples were extracted and diluted with distilled water. Standards were made of BSA stock solution, and 25 µL of each standard and protein dilution were put into a 96-well plate (Bio-Rad). The bicinchoninic acid (BCA) protein assay kit (Pierce BCA Protein Assay kit; Thermo Scientific) was made up according to the manufacturer's instructions, and 200 µL was added to each well. The amount of protein in each well was estimated using absorbance readings from a SPECTROstar Nano plate Reader (BMG Lab-Tech). Protein samples (20 µg) were loaded onto a 10% Mini-PROTEAN TGX precast gel (Bio-Rad) alongside 5 µl normal molecular weight ladder (Precision Plus Protein Ladder; Bio-Rad) at 200 V for 45 min and then transferred to an iBlot 2 PVDF mini

stack membrane (Invitrogen) using an iBLot 2 (ThermoFisher Scientific). The membranes were blocked in 3% BSA in TBST for 1 h. Membranes were transferred to primary antibody solutions (rabbit anti-MMP7 antibody #NBP1-99123 (1.28 mg/mL) Novus Biologicals) diluted in 3% BSA in TBST and left on a spinner for 24 h at 4°C. Membranes were then rinsed three times for 5 min each in TBST and transferred to a secondary antibody solution (polyclonal goat anti-rabbit IgG horseradish peroxidase (HRP)-linked antibody #HAF1008 (0.1 mg/mL) R&D Systems) in 3% BSA for 1 h at 4°C. Then, membranes were rinsed in TBST and immediately imaged using a Gel Doc (Syngene G:BOX-Chemi-QRX using enhanced chemiluminescence (Clarity™ Western ECL Substrate: Bio-Rad).

*RNA sequencing and analysis*
Uterine samples were snap frozen in liquid nitrogen and stored at -80°C. Tissue was sampled from non-gravid and gravid uteri from pregnant females based on the number of days after the removal of pouch young. The stages included pre-ovulation (sample collected from the uterus contralateral to the post-partum uterus within 24 hours of birth; n=2), pre-attachment (day 14; gravid n=3, non-gravid n=2), early attachment (day 20; gravid n=3, non-gravid n=2), late attachment (day 25; gravid n=3, non-gravid n=3) and post-partum (sample collected within 24 hours of birth; n=2). RNA was extracted from snap frozen tissue by mechanical disruption with a Qiagen Tissue Ruptor and an RNeasy mini kit (Qiagen) following standard protocols. RNA quality and quantity was measured and RNA samples with a with an RNA Integrity Number (RIN) greater than 8 was used, which is high enough quality for all downstream applications. Illumina sequencing libraries were generated by the Ramaciotti Center for Genomics (UNSW Sydney, Australia) using the TruSeq Stranded mRNA prep and sequenced with NovaSeq S1 100bp sequencing. A mean of 45.6 million reads were sequenced per sample. Raw reads were aligned to the tammar wallaby genome v3.0 using hisat2 and reads were counted using htseq-count. A mean mapping rate of 92% was achieved. Hierarchical clustering and principal component analysis (PCA) was performed in the R *stats* package (65) to confirm that there were general patterns in gene expression between treatment groups. We compared differential gene expression analysis between non-gravid PP, d14, d20, d25 and pre-ovulation (PO), gravid d14, d20, d25 uteri using DeSeq2 default parameters (66). Read counts were normalized and presented as adjusted P values for multiple comparisons (Appendix Table S1-S8). Differential expression was called with a false discovery rate of 0.05. Gene ontology was analyzed using GOrilla and visualized with REVIGO (Appendix Table S9-S21, 67).

#### Supplementary results

*Western blotting*
Western blots of whole uterine tissue (20 µg) from tammar wallaby uterine tissue showed bands at the expected size for MMP7 of 40kDa. MMP7 has a full length of 50 kDa with proteolytic fragments (68). In the present study, bands for MMP7 were visible around 50, 37 and 15 kDa for all animals.

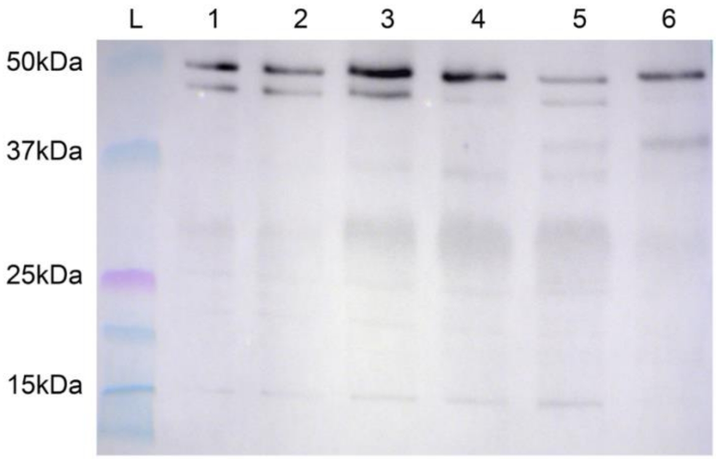


***Fig. 8.*** Western blot of protein extracted from tammar wallaby endometrium (20 µg protein loaded per well) at varying stages of pregnancy. Bands for MMP7 occur at 50, 37 and 15 kDa. L, ladder, ***1.*** Day 14 gravid ***2.*** Day 14 non-gravid ***3***. Day 20 gravid ***4.*** Day 20 non-gravid ***5.*** Day 25 gravid ***6.*** Day 25 non-gravid.
